# A pivotal genetic program controlled by thyroid hormone during the maturation of GABAergic neurons in mice

**DOI:** 10.1101/574202

**Authors:** Sabine Richard, Romain Guyot, Martin Rey-Millet, Margaux Prieux, Suzy Markossian, Denise Aubert, Frédéric Flamant

## Abstract

In mammals, brain development is critically dependent on proper thyroid hormone signaling, via the TRα1 nuclear receptor. However, the downstream mechanisms by which TRα1 impacts brain development are currently unknown, notably because this receptor is expressed ubiquitously from early stages of development. In order to better define the function of TRα1 in the developing brain, we used mouse genetics to induce the expression of a dominant-negative mutation of the receptor specifically in GABAergic neurons, the main inhibitory neurons in the brain, which were previously identified as sensitive to hypothyroidism. This triggered post-natal epileptic seizures, reflecting a profound impairment of GABAergic neuron maturation in different brain areas. Analysis of transcriptome and TRα1 cistrome also allowed us to identify a small set of genes, the transcription of which is upregulated by TRα1 in GABAergic neurons during post-natal maturation of the striatum and which probably play an important role during neurodevelopment. Thus, our results point to GABAergic neurons as direct targets of thyroid hormone during brain development and suggest that many defects seen in hypothyroid brains may be secondary to GABAergic neuron malfunction.

## Introduction

Thyroid hormones (TH, including thyroxine, or T4, and 3,3’,5-triiodo-L-thyronine, or T3, its active metabolite) exert a broad influence on neurodevelopment. If untreated soon after birth, congenital hypothyroidism, i.e. early TH deficiency, affects brain development and a number of cognitive functions (1). Severe cases display mental retardation, autism spectrum disorders (ASD) and epilepsy (2). TH mainly acts by binding to nuclear receptors called TRα1, TRβ1 and TRβ2, which are encoded by the *THRA* and *THRB* genes (*Thra* and *Thrb* in mice, formerly *TRα* and *TRβ*). These receptors form heterodimers with other nuclear receptors, the Retinoid-X-Receptors (RXRs), and bind chromatin at specific locations (thyroid hormone response elements), acting as TH-dependent transcription activators of neighboring genes. In the developing brain, the predominant type of receptor is TRα1 (3). Accordingly, *THRA* germline mutations, which have currently been reported in only 45 patients, cause a syndrome resembling congenital hypothyroidism, with mental retardation and a high occurrence of ASD and epilepsy (4).

Cellular alterations caused by early TH deficiency or germline mutations have been extensively studied in rodents. Many neurodevelopmental processes depend on proper TH signaling, and virtually all glial and neuronal cell populations are affected by TH deficiency (5, 6). However, in the mouse cerebellum, we have previously found that, although *Thra* expression is ubiquitous in this brain region, only a subset of cell types displayed a direct, cell-autonomous, response to TH (7). More specifically, Cre/loxP technology, used to express a dominant-negative variant of TRα1, has provided genetic evidence that the cell-autonomous influence of TRα1 is limited to astrocytes and GABAergic neurons (7). By altering the differentiation of GABAergic neurons, TRα1^L400R^ prevented the secretion of several growth factors and neurotrophins. This indirectly altered the proliferation and differentiation of granule cells and oligodendrocytes (8). Therefore GABAergic neurons occupy a pivotal position during cerebellum development, amplifying the initial TH signal. This allows TH to synchronize cellular interactions and the maturation of neuronal networks during the first post-natal weeks (9).

In the present study, we used the same strategy to ask whether the influence of TH on GABAergic neurons can be generalized to other brain areas. We found that blocking TH response by expressing TRα1^L400R^ in the entire GABAergic lineage from early developmental stages had dramatic neurodevelopmental consequences in mice, causing lethal epileptic seizures. Development of the GABAergic system was greatly altered. Genome-wide analyses allowed us to pinpoint the genetic defects induced by the mutation, and to identify a small set of genes activated by TH in GABAergic neurons. These genes are likely to play a key role in the neurodevelopmental function of TH.

## Results

### Mouse models

We generated new mouse models by combining existing and novel “floxed” *Thra* alleles with the *Gad2Cre* transgene (Fig. 1a). This transgene drives the expression of Cre recombinase in all GABAergic neurons and their progenitors from an early prenatal stage (around E12.5 (10)). In the context of the modified *Thra* alleles used in the present study, Cre recombinase eliminates a transcriptional stop cassette and triggers the expression of TRα1 variants. The *Thra*^*AMI*^ allele (formerly named *TRα*^*AMI*^ (11)) encodes TRα1^L400R^ (Fig. 1b), which exerts a permanent transcriptional repression on target genes, even in the presence of TH. This is due to its inability to recruit transcription coactivators, which normally interact with the C-terminal helix (AA 398-407) of TRα1. This allelic design, which eliminates alternate splicing, increases the expression of the mutant receptor, over that of the wild-type receptor. This ensures a complete inhibition of TH response in heterozygous cells (12). Like complete deprivation of TH (13), ubiquitous expression of TRα1^L400R^ results in the death of heterozygous mice 2-3 weeks after birth (11). The second modified *Thra* allele, named *Thra*^*Slox*^, differs from *Thra*^*AMI*^ only by an additional frameshift mutation, which eliminates the C-terminal helix of TRα1 (12). This allele encodes TRα1^E395fs401X^ (Fig. 1b), which is nearly identical to a pathological variant found in a patient (14) and is expected to be functionally equivalent to TRα1^L400R^. The third modified *Thra* allele is *Thra*^*TAG*^ a novel construct, which encodes a functional receptor, TRα1^TAG^, with a fragment of protein G at its N-terminus (15). This tag has a high affinity for IgGs, which makes it suitable to address chromatin occupancy (16). Double heterozygous mice, combining the presence of *Gad2Cre* and of a modified *Thra* allele, express TRα1^L400R^, TRα1^E395fs401X^ or TRα1^TAG^ in the GABAergic cell lineage only. They will be respectively designated as *Thra*^*AMI/gn*^ *Thra*^*Slox/gn*^ and *Thra*^*TAG/gn*^ in the following. In all phenotyping experiments, littermates carrying only *Thra*^*AMI*^, *Thra*^*Slox*^ or *Gad2Cre* were used as controls.

**Figure 1:**
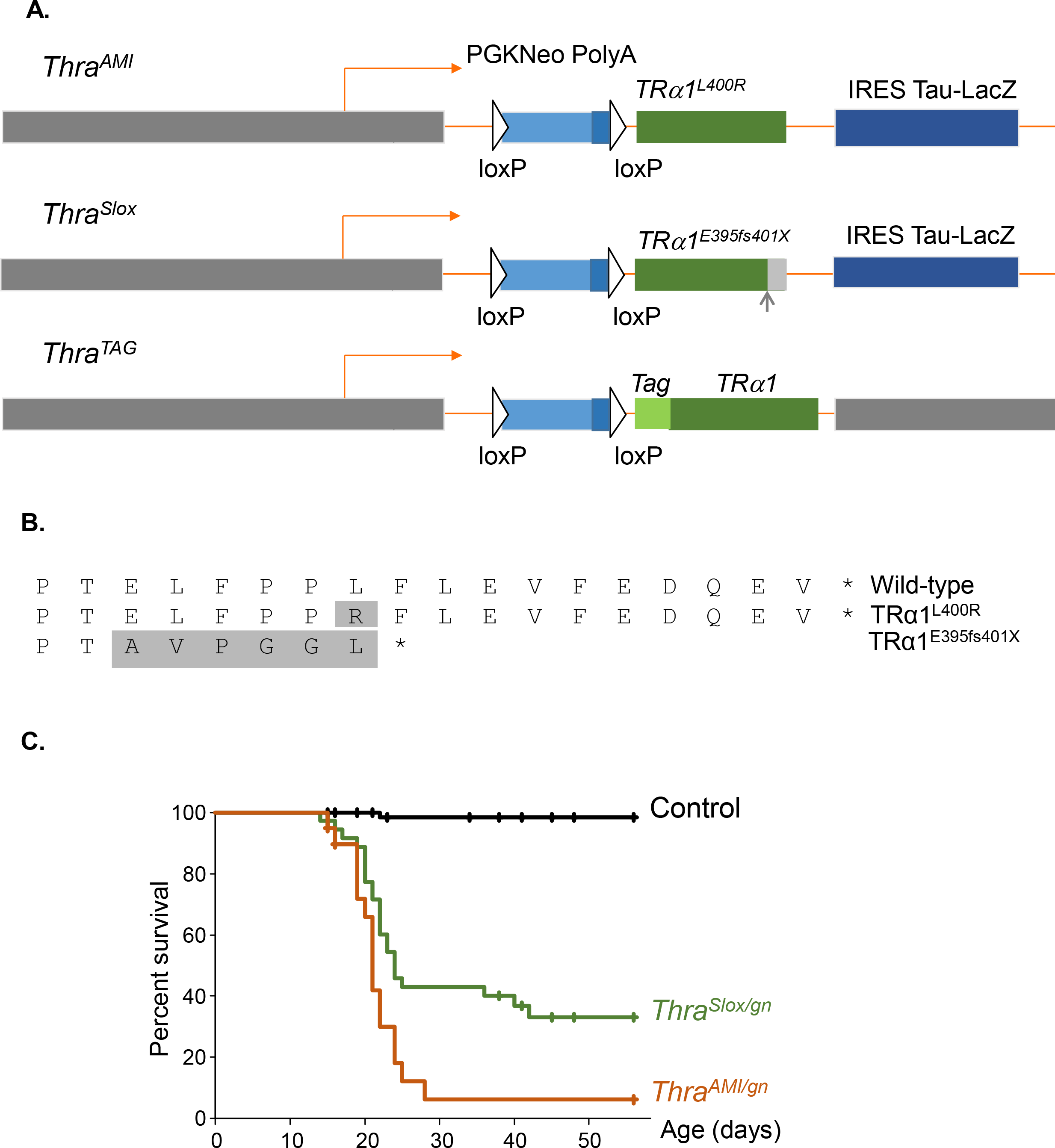
*Thra* alleles and survival curves. **A.** Schematic representation of *Thra* alleles in *Thra*^*AMI*^, *Thra*^*Slox*^ and *Thra*^*TAG*^ mice. In all 3 alleles, the coding sequence is preceded with a floxed stop cassette (PGKNeo PolyA). The intronless structure eliminates alternate splicing and internal promoter, and thus prevents the production of TRα2, TRΔα1 and TRΔα2 non-receptor protein. The dispensable IRES Tau-lacZ reporter part was not included in the *Thra*^*TAG*^ construct. **B.** C-terminal amino-acid sequence of *Thra* gene products used in the present study, starting from AA393. Shaded amino-acids differ from wild-type TRα1. *Thra*^*AMI*^ mutation results in a single amino-acid substitution within TRα1 helix 12. *Thra*^*Slox*^ mutation is a deletion resulting in a +1 frameshift, leading to elimination of helix 12, as in several RTHα patients. **C.** Survival curves of mice expressing a mutated TRα1 in GABAergic neurons (green lines) and of control littermates (black line).

### Post-natal lethality caused by Thra mutations in GABAergic neurons

Born at the expected frequency, *Thra*^*AMI/gn*^ mice did not usually survive beyond the third post-natal week (Fig. 1c). Video recording of litters in their home cage indicated that most mice started to display epileptic seizures a few days before death. Occasionally, sudden death was observed at the end of a seizure. In most cases, seizures impeded maternal care and this likely precipitated the death of the pups (Suppl. movie S1 and movie S2). Although lethality was also observed in *Thra*^*Slox/gn*^ mice (Fig. 1c), about one third of these mice survived into adulthood. Adult *Thra*^*Slox/gn*^ mice did not display any obvious epileptic seizure anymore. However, their locomotor behavior was significantly altered, as evidenced in an open-field test (Suppl. info. fig. S1). These observations show that, although the two mutations are expected to be equivalent, TRα1^E395fs401X^ is less detrimental than TRα1^L400R^.

### A global impairment in the differentiation of GABAergic neurons

In order to label neurons of the GABAergic lineage in a generic way, we combined the *Thra*^*AMI/gn*^ and *Gad2Cre* alleles with the *Rosa-tdTomato* transgene, which enabled to trace the cells in which *Cre/loxP* recombination had taken place. The density of tdTomato+ cells was not reduced in *Thra*^*AMI/gn*^/*Rosa-tdTomato* mice, arguing against a possible alteration in the proliferation, migration or survival of GABAergic neuron progenitors. In the hippocampus, notably in the dentate gyrus, the number of tdTomato+ cells was even increased (Fig. 2).

**Figure 2:**
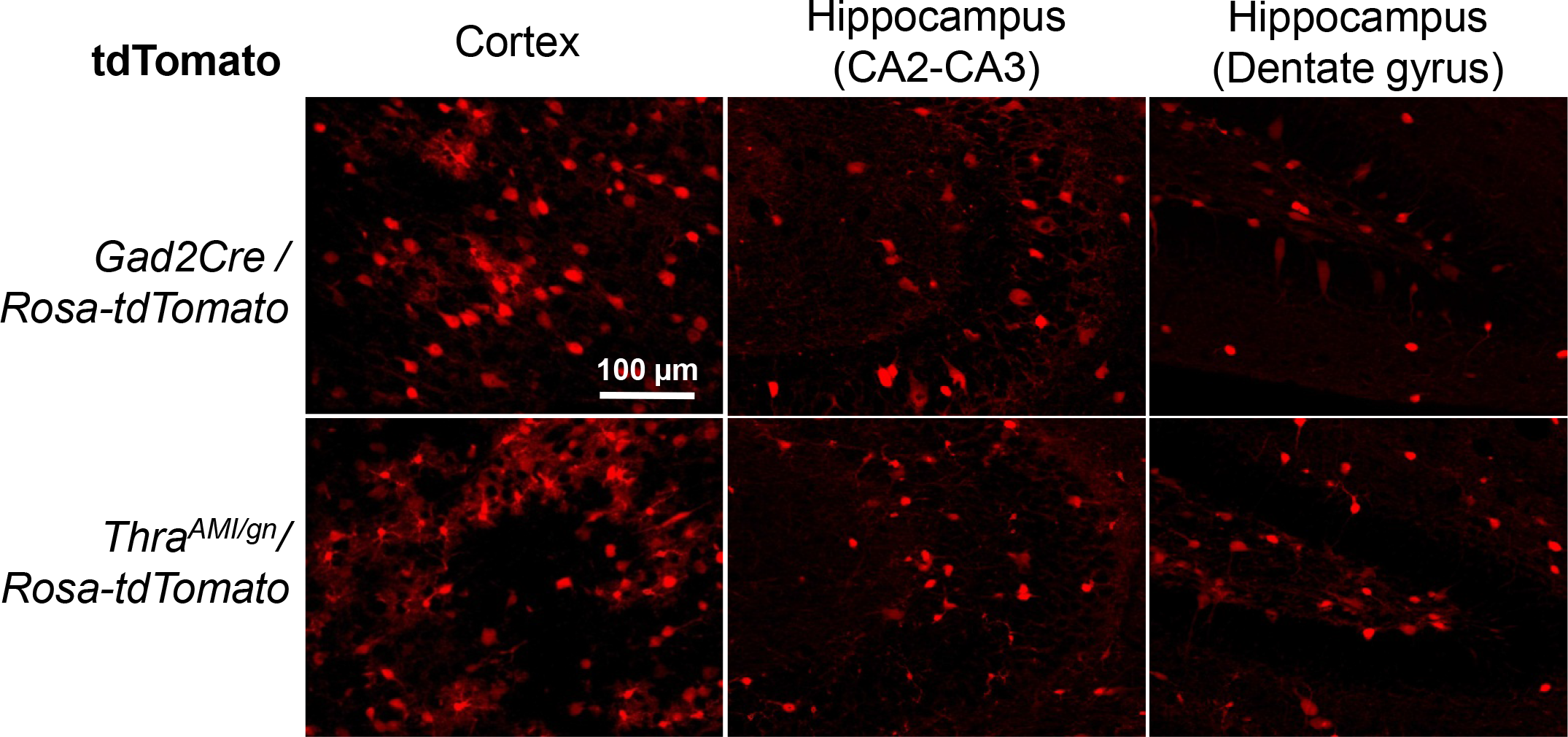
Expression of the tdTomato fluorescent protein in *TRα*^*AMI/gn*^/*Rosa-tdTomato* and control littermates at PND14. The relative density of fluorescent cells in mutant mice was unchanged in the cortex (1.00±0.09, *n*=6 in controls *vs* 1.08±0.11, *n*=6 in mutant mice; *ns*) and slightly increased in the CA2-CA3 area (1.00±0.05, *n*=6, in controls *vs* 1.20±0.07, *n*=5 in mutant mice; *p* <0.05) and dentate gyrus (1.00±0.06, *n*=6 in controls *vs* 1.47±0.10 *n*=6 in mutant mice; *p* <0.05) of the hippocampus.

We used immunohistochemistry to detect alterations of GABAergic neuron differentiation at post-natal day 14 (PND14) in various brain areas (Suppl. info. table S1). Parvalbumin (PV) immunostaining, which labels major populations of GABAergic neurons in several brain areas (17), revealed a defect in Purkinje cell arborization, and a deficit in basket and stellate GABAergic interneurons in *Thra*^*AMI/gn*^ cerebellum as expected from previous data (12, 18). A drastic reduction in the density of PV+ neurons was also visible in the hippocampus, cortex and striatum of *Thra*^*AMI/gn*^ mice (Fig. 3a). We also used antibodies directed against neuropeptide Y (NPY), calretinin (CR) and somatostatin (SST) to label other key populations of GABAergic neurons (19). All GABAergic neuron subtypes investigated were affected, but in a rather complex pattern. The density of NPY+ neurons was significantly reduced in the cortex, but not in the striatum, where NPY immunoreactivity of the fibers increased nevertheless (Suppl. info. fig. S2). The density of CR+ neurons was increased in the cortex and CA2-CA3 area of the hippocampus. In the hippocampal dentate gyrus, where CR+ neurons are normally concentrated in the granular cell layer, the limits of this layer were ill-defined and CR+ neurons spread into the molecular and polymorph layers (Fig. 3b). The density of SST+ neurons was also augmented in hippocampal dentate gyrus, but not significantly altered in the cortex or striatum (Fig. 3c).

**Figure 3:**
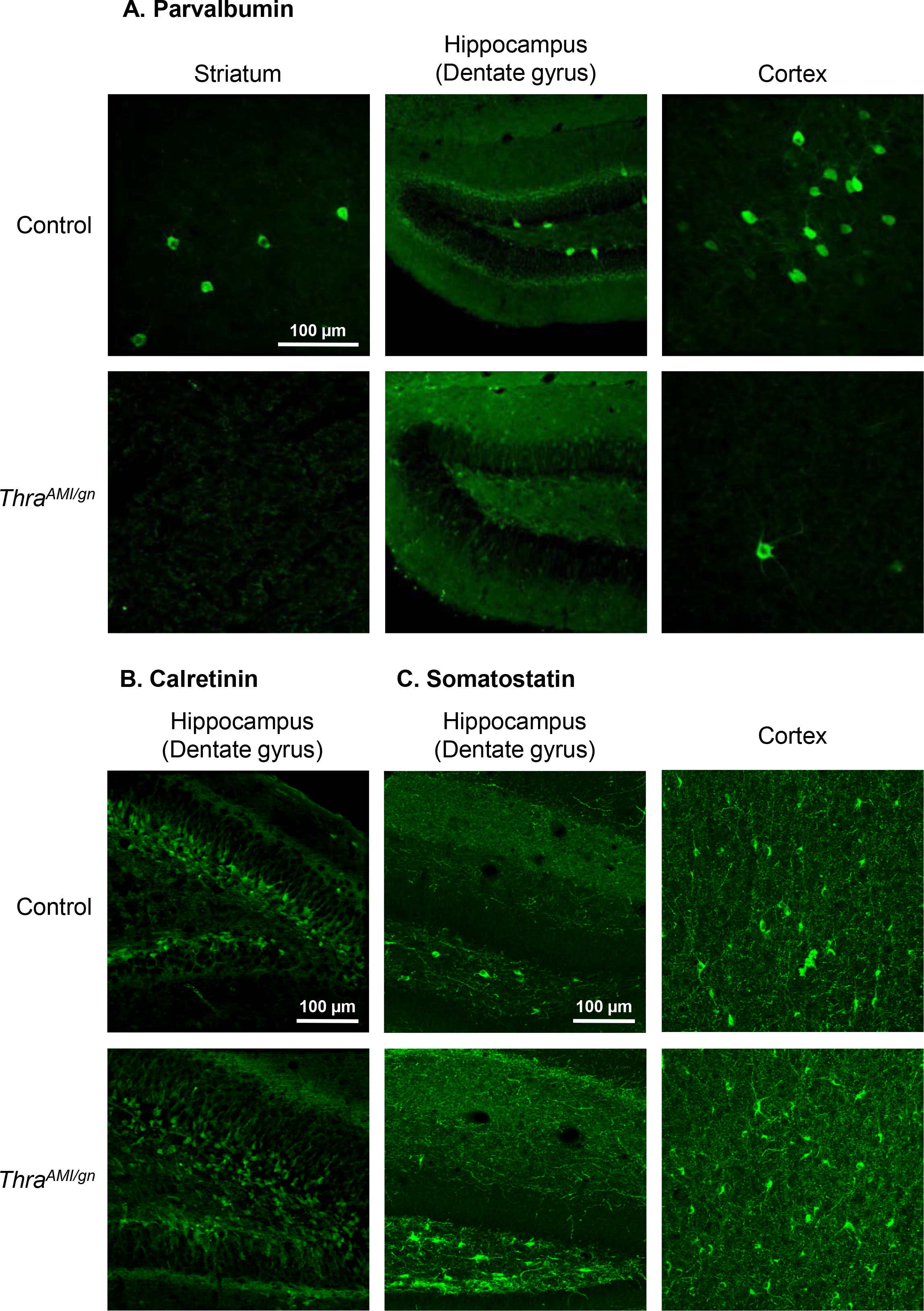
Immunohistochemistry for parvalbumin (A), calretinin (B) and somatostatin (C) in PND14 *Thra*^*AMI/gn*^ and control mouse pups in selected brain regions.

Most of the above-mentioned immunohistochemistry experiments were also carried out in *Thra*^*Slox/gn*^ mice at PND14. In all instances, the defects observed in GABAergic neuron populations were the same as in *Thra*^*AMI/gn*^ mice (Details in Suppl. info. table S2 and fig. S3). In the surviving *Thra*^*Slox/gn*^ adult mice, PV and NPY immunohistochemistry indicated that the differentiation of neurons expressing these markers was not only delayed, but permanently impaired (Suppl. info. fig. S4 and table S2). Taken together, these data indicate that expressing a mutant TRα1 alters the terminal differentiation of GABAergic neurons, reducing the numbers of PV+ and NPY+ cells in some brain areas, while favoring an expansion of the SST+ and CR+ cell populations. In the hippocampus, this is accompanied by subtle morphological alterations.

### Changes in gene expression caused by TRα1^L400R^ in GABAergic neurons of the striatum

Because the impairment in GABAergic neuron differentiation does not appear to be restricted to a specific brain area or a specific neuronal GABAergic subpopulation, we hypothesized that a general mechanism might underlie the involvement of TH signaling in the terminal maturation of GABAergic neurons. In order to decipher this mechanism, and identify the TRα1 target genes in GABAergic neurons, we focused our investigation on the striatum, where the high abundance of GABAergic neurons, mainly spiny projection neurons, facilitates the interpretation of gene expression analyses. We compared the transcriptome of the striatum in *Thra*^*AMI/gn*^ and control littermates at two different post-natal stages, PND7 and PND14, using the Ion AmpliSeq Transcriptome Mouse Gene Expression Assay. Differential gene expression analysis (Deseq2 software, adjusted p-value inf. 0.05; FDR=0.05; fold-change >2 or <0.5; mean normalized expression >10) pointed out a set of 126 genes whose expression was deregulated at PND7 in *Thra*^*AMI/gn*^ mouse striatum, compared to control mice (98 down-regulated; 28-up regulated). At PND14, this number raised to 215 genes (144 down-regulated; 71 up-regulated). A two-factor analysis increased the statistical power for genes deregulated at both stages, and added 8 genes to these lists. Overall, 260 genes were found to be deregulated in the striatum of *Thra*^*AMI/gn*^ mice at either stage. Hierarchical clustering analysis of these data mainly revealed a sharp transition between PND7 and PND14 in the striatum of control mice, which did not take place in *Thra*^*AMI/gn*^ mice (Fig. 4).

**Figure 4:**
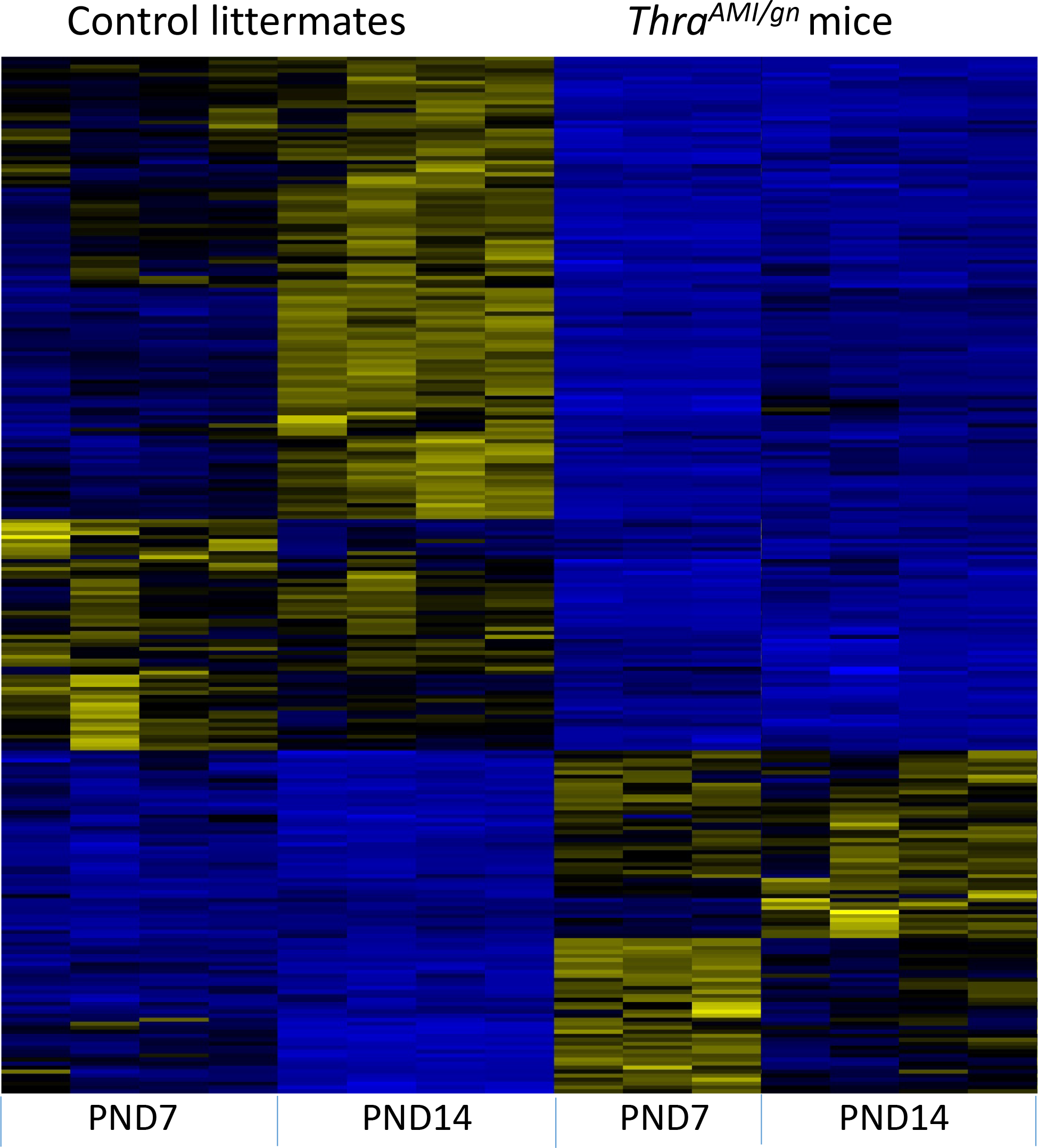
Hierarchical clustering analysis of differentially expressed transcripts in the striatum of *Thra*^*AMI/gn*^ mice and control littermates at PND7 and PND14. The analysis is restricted to 260 genes for which the fold-change is >2 or <0.5 (adjusted *p*-value < 0.05) for at least one developmental stage. High expression is in yellow, low expression is in blue, average in black. Note that the changes in gene expression between PND7 and PND7 are more noteworthy in control than in mutant mice, suggesting that a maturation process is blunted by the mutation.

These differences in gene expression, as measured by Ampliseq, may have two origins. They can reflect deregulations of the expression of TRα1 target genes in GABAergic neurons, but they can also reflect various indirect consequences of these deregulations, such as a change in the composition of the cell population. In order to pinpoint the TRα1 target genes within the set of differentially expressed genes, we crossed the above results with a dataset obtained in wild-type mice at PND14, comparing different hormonal statuses. We assumed that the expression of TRα1 target genes should be quickly modified by changes in TH levels, while indirect consequences should be much slower. In the dataset, 181 genes were found to be deregulated in the striatum of hypothyroid mice, while 86 genes responded to a 2-day TH treatment of hypothyroid mice (Suppl. Dataset S1). In this dataset however, the response of GABAergic neurons to TH cannot be distinguished from the response of other cell types present in the striatum, which also express TRα1. The overlap between the two datasets pointed out a set of 38 TH-activated genes whose expression pattern suggested a direct regulation by TRα1 in GABAergic neurons, while only 1 gene displayed an expression pattern suggestive of a negative regulation by T3-bound TRα1 (Fig. 5a).

**Figure 5:**
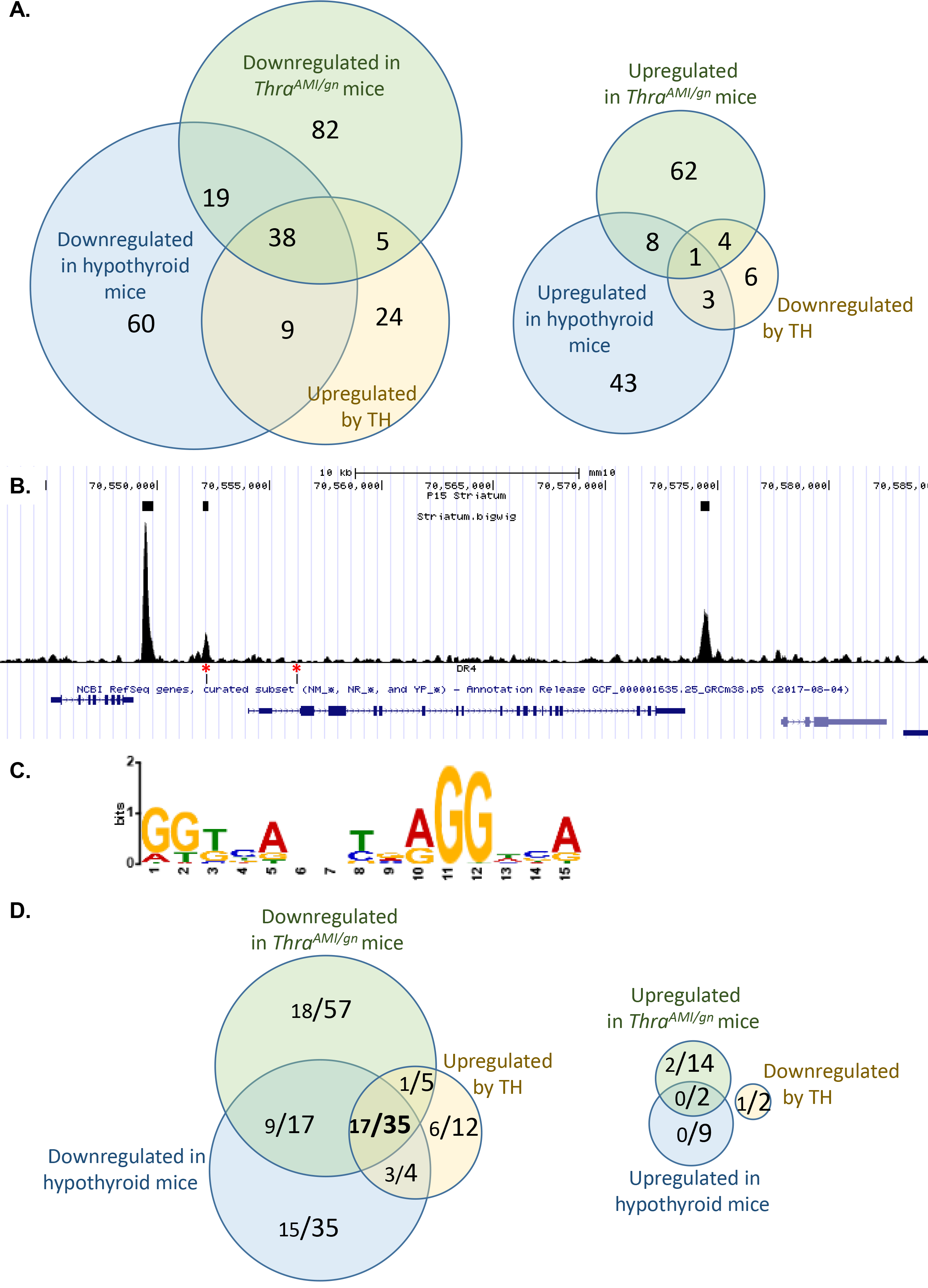
Identifying a core set of TRα1 target genes in GABAergic neurons of the striatum by combining RNAseq and Chip-Seq analyses. **A.** RNAseq identifies a set of 38 genes, the expression pattern of which is fully consistent with a positive regulation by TRα1, and only 1 gene which has the opposite expression pattern. **B.** Extract of the *Mus musculus* genome browser, around the *Hr* gene, a well-characterized TRα1 target gene. The 3 upper boxes indicate TRBSs identified as significant by the MACS2 algorithm. Note that a DR4-like element (lower track, red asterisks), as defined below, is found in only one of the 3 peaks. **C.** Consensus sequence found in TRBSs identified by *de novo* motif search is close to the DR4 consensus (5’AGGTCANNNNAGGTCA3’). **D-E.** Combinations of RNAseq and Chip-Seq data. In the Venn diagrams, each fraction gives the number of genes with a proximal TRBS (<30 kb for transcription start site, large lettering) and, among these genes, those in which a DR4 element was identified (small lettering). A set of 35 genes fulfil the criteria for being considered as genuine TRα1 target genes: they are down-regulated in hypothyroid and mutant mice, and up-regulated after TH treatment of hypothyroid mice. For 17 of these genes, the TRBS contains a recognizable DR4-like element.

### Identification of direct TRα1 target genes in GABAergic neurons of the striatum

To complete the identification of TRα1 target genes in striatal GABAergic neurons, we analyzed chromatin occupancy by TRα1 at a genome-wide scale by ChIPseq. Using *Thra*^*TAG/gn*^, we precipitated the DNA/protein complexes, which contain the tagged TRα1 from the whole striatum, to address chromatin occupancy in GABAergic neurons only. This experiment revealed 7,484 sites occupied by TRα1^TAG^ (thyroid hormone receptor binding sites = TRBS) in the genome (Fig. 5b). In agreement with our previous study (16) *de novo* motif discovery (http://meme-suite.org/tools/meme-chip) and enrichment analysis revealed a single consensus sequence for the binding of TRα1/RXR heterodimers. The sequence is the so-called DR4 element (Fig. 5c). Assuming that proximity (<30kb) between the TRα1^TAG^ binding site and the transcription start site is sufficient for direct transcriptional regulation by TRα1 would lead to consider a large fraction of the genes as putative target genes (3,979/23,931 annotated genes in the mouse genome mm10 version; 16.6%). Among the 38 genes which are sensitive to TRα1^L400R^ expression and hypothyroidism and responsive to TH in hypothyroid mice, genes with a proximal TRBS were overrepresented (35/38: 92%; enrichment of 5.5 compared to the whole set of annotated genes). Interestingly, this enrichment was more striking if we considered only the TRBSs where a DR4 element was identified (3,813/7,484; 51%): DR4 elements were present within 30 kb of 7.5% of annotated genes (1,786/23,931), and of 45% of the putative TRα1 target genes identified in the present study (17/38)(Fig. 5d). The same reasoning leads to the conclusion that the transcription of the genes which are down-regulated after T3 treatment is not regulated by chromatin-bound TRα1 (Fig. 5e). Overall, the ChipSeq dataset suggests that a large fraction of the TRBSs does not reflect the binding of TRα1/RXR heterodimers to DR4 elements, but corresponds to other modes of chromatin association, which do not necessarily promote TH-mediated transactivation. However, as we cannot rule out that other types of response elements are also used, this analysis leaves us with 35 genes, which meet all the criteria for being considered as TRα1 direct target genes. Although some of these genes are known to have a neurodevelopmental function, they do not fall into a specific ontological category (Suppl. info. table S3). This implies that TH promotes GABAergic neuron maturation by simultaneously acting on different cell compartments and cellular pathways.

## Discussion

Using two mouse models expressing mutant forms of TRα1 specifically in GABAergic neurons, we present novel evidence showing that TH bound to TRα1 promotes the terminal differentiation and maturation of GABAergic neurons. As we used here GABAergic-specific somatic mutations, we can ascertain that the observed defects are cell-autonomous consequences of impaired TH signaling. We found that the defect in GABAergic differentiation is not restricted to a specific brain area, nor to a specific subtype of GABAergic neurons, as several subtypes of both projecting neurons and interneurons were affected by TRα1 mutations. This suggests that, although GABAergic neurons of different brain areas have different embryonal origins (20, 21), they share a common pathway of maturation that depends on TH/TRα1 signaling. These results extend previous findings on the role of TH in GABAergic neurons in the cerebellum (18, 22), striatum (23), cortex (24), hippocampus (25) and hypothalamus (26). In many respects, neurodevelopmental damage caused by TRα1^L400R^ and TRα1^E395fs401X^ appears to be more dramatic than that reported for the TRα1^R384C^ mutation (24), which is impaired in its affinity for TH, but possesses a residual capacity to transactivate gene expression (27).

The present data indicate that TRα1 plays a major role in the terminal steps of differentiation of several categories of GABAergic neurons. This is most obvious for PV+ interneurons, which almost disappear from several brain areas in *Thra*^*AMI/gn*^ mice. However, their progenitors appear to be present, as evidenced by the use of tdTomato as a reporter for cells of the GABAergic lineage. The fate of these progenitors is unclear. One hypothesis is that they commit to a different GABAergic lineage. This could notably explain the excess of SST+ cells in the hippocampus, since PV+ and SST+ cortical interneurons share the same precursors (28, 29). However, the results obtained in the cortex do not support such hypothesis. An alternative hypothesis would be that the effects observed in different categories of GABAergic neurons are secondary to the near disappearance of PV+ interneurons. Indeed, many defects caused by hypothyroidism in the brain are secondary to a defect in neurotrophin secretion in the microenvironment (30–33).

Our genome-wide search pinpointed a small set of genes fulfilling the criteria which lead us to consider them as genuine TRα1 target genes in GABAergic neurons: (1) the mRNA level of these genes is TH responsive, (2) it is decreased in the striatum of *Thra*^*AMI/gn*^ mice and (3) TRα1 occupies a chromatin binding site located at a limited distance of their transcription start site. The last criterion is important, as it helps to differentiate between direct and indirect influence of TRα1 on gene regulation. It also indicates that the negative regulation of gene expression, also exerted by TH, is not directly mediated by chromatin-bound TRα1, but perhaps an indirect consequence of the upregulation of a transcription inhibitor. To our knowledge, this study is the first to address chromatin occupancy by TRα1 *in vivo*. Importantly, it does so using a genetic strategy which enables to selectively identify GABAergic neuron-specific TRα1 binding to DNA within a heterogeneous tissue. Interestingly, the large number of chromatin binding sites that we have identified (7,484 TRBS) contrasts with the small set of 35 genes that we have identified as being directly regulated by TRα1 in GABAergic neurons. As we have used stringent statistical thresholds, we have probably overlooked some genuine TRα1 genes. For example, *Klf9* is a known target gene (34) which is not present in our list, due to a modest downregulation in hypothyroid striatum. However, even with liberal statistical thresholds, the number of presumptive target genes will not exceed 100, a number which is still small compared to the number of genes with a proximal TRBS. Such a contrast has previously been observed in other systems (16, 35, 36) and suggests that only a small fraction of the TRBSs are involved in TH mediated transactivation. Further investigations will be required to better define these active TRBSs, and better establish the correspondence between chromatin occupancy and transcriptional regulation by TRα1.

Most of the TRα1 target genes identified in the striatum have already been identified as being sensitive to the local TH level in various brain areas and at different developmental stages (see supplementary Table S1 in ref. 34). This reinforces the hypothesis that they belong to a common genetic program which is regulated by TH, via TRα1, and which promotes the proper maturation of several categories of GABAergic neurons. Although their function in neurons is for a large part unknown, these genes can be grouped according to the putative function of their protein products: *Shh* and *Fgf16* encode secreted proteins, which play major roles in cellular interactions. *Sema7a* and *Nrtn* are involved in axon growth and pathfinding. Others are likely to define the electrophysiological properties of the neurons by encoding ion channels *(Kctd17)*, transporters of small metabolites *(Slc22a3, Slc26a10)* or modulators of synaptic activity (*Nrgn*, *Lynx1*).

Overall, the broad influence of TRα1 mutations on GABAergic neuron differentiation and maturation is expected to greatly and permanently impair brain function, notably in the cortex, where a subtle equilibrium between different GABAergic neuron subtypes is necessary for normal development and plasticity (37). In the mouse models presented here, epileptic seizures appear to be a main cause of mortality, which sheds light on the cause of the lethality that had been previously observed in mice with different *Thra* mutations (11, 27, 38, 39), as well as in mice suffering from complete TH deprivation (13, 40). This phenotype is highly relevant to human pathology, as a history of epilepsy has been reported for several of the rare patients with a *THRA* mutation (41). Autism spectrum disorders (ASD), whose comorbidity with epilepsy is well documented, have also been reported in these patients (42, 43). It is likely that these pathological traits are also due to a defect in GABAergic neurons maturation and our data suggest that these patients might benefit from a treatment with GABA receptor agonists.

## Materials and methods

### Mouse models and treatments

All experiments were carried out in accordance with the European Community Council Directive of September 22, 2010 (2010/63/EU) regarding the protection of animals used for experimental and other scientific purposes. The research project was approved by a local animal care and use committee and subsequently authorized by the French Ministry of Research. Mice were bred and maintained at the Plateau de Biologie Expérimentale de la Souris (SFR BioSciences Gerland - Lyon Sud, France).

In order to target the expression of mutated forms of TRα1 specifically in GABAergic neurons, we used the Cre/loxP system with *Gad2Cre* mice, which carry a Cre coding cassette immediately after the translation stop codon of the *Gad2* gene. In *Gad2Cre* mice, Cre recombinase expression is almost entirely restricted to GABAergic neurons and includes almost all GABAergic neurons (10). *Gad2Cre* mice were crossed with *Thra*^*AMI*/+^ (11) or *Thra*^*Slox/+*^ (12) mice, which carry mutated versions of the *Thra* gene preceded with a floxed STOP cassette (Fig. 1). When indicated, a *Rosa26tdTomato* reporter transgene (also known as *Ai9*) was also included (44). Finally, in order to address chromatin occupancy by TRα1 specifically in GABAergic neurons, we generated *ad hoc Thra*^*TAG/gn*^ transgenic mice, which express a “tagged” receptor, TRα1^TAG^, only in GABAergic neurons (Fig. 1). In order to compare animals with different thyroid statuses, TH deficiency was induced in pups by giving a diet containing propylthiouracil (TD95125; Harlan Teklad) to dams from post-natal day 0 (PND0) to post-natal day 14 (PND 14). In half of hypothyroid pups, TH levels were restored by two intraperitoneal injections of a mixture of T4 (2g/kg) and T3 (0.2g/kg) given at PND12 and PND13.

### Brain collection

For neuroanatomy experiments, each mouse was given a lethal intraperitoneal injection (6 mL/kg) of a mixture of ketamine (33 mg/mL) and xylazine (6.7 mg/mL). The thorax was opened and each mouse was perfused with 4% paraformaldehyde in 0.1M phosphate buffer at room temperature. Each brain was dissected out, immersed in fixative at 4°C for 3 hours and then in phosphate buffered saline (PBS) at 4°C until sectioning. Coronal sections (50 μm) were cut with the aid of a vibrating microtome (Integraslice 7550 SPDS, Campden Instruments, Loughborough, UK), in PBS at room temperature. Brain sections were stored at −20°C in cryoprotectant (30% ethylene glycol and 20% glycerol in 10 mM PBS) prior to immunohistochemistry. For transcriptome analyses, pups were killed by decapitation. The striatum was dissected out, snap frozen in either dry ice or liquid nitrogen and stored at −80°C prior to analysis. For Chip-Seq experiments, the tissue was dissociated immediately after dissection and DNA-protein complexes were cross-linked by incubation in 1% formaldehyde for 20 min.

### Immunohistochemistry

Immunohistochemistry was performed on free-floating sections. The following primary antibodies were used: mouse anti-parvalbumin (Sigma P3088, 1:2000), rabbit anti-NPY (Sigma N6528, 1:5000), goat anti-somatostatin (Clinisciences D20 sc-7819, 1:500) and mouse anti-calretinin (Swant 6B3, 1:1000). Secondary antibodies were made in donkey (anti-rabbit, anti-goat and anti-mouse DyLight 488, ThermoFisher Scientific) and used at a 1:1000 dilution. Antibodies were diluted in PBS with 0.2% Triton X-100, 1% normal donkey serum, 1% cold water fish skin gelatin and 1% dimethyl sulfoxide. Non-specific binding sites were blocked by incubating sections for 1 h in PBS with 10% normal donkey serum, 1% cold water fish skin gelatin and 0.2% Triton X-100. Primary antibody incubation was carried out overnight at 4°C and sections were washed in PBS with 1% normal donkey serum and 1% cold water fish skin gelatin. Sections were further incubated for 10 min at room temperature with DAPI (4’,6-diamidino-2-phenylindole, 1:5000, Sigma). Incubation with the secondary antibodies lasted for 3h at room temperature. Sections were mounted in FluoroshieldTM (Sigma), coverslipped and imaged using an inverted confocal microscope (Zeiss LSM 780).

### Transcriptome analysis

RNA was extracted from the striatum using a Macherey-Nagel NucleoSpin RNA II kit. RNAseq was used to compare gene expression levels between control mice (n=4), hypothyroid mice (n=5) and hypothyroid mice treated for 48 hours with T4 and T3 (n=5). cDNA libraries were prepared using the total RNA SENSE kit (Lexogen, Vienna Austria) and analyzed on an Ion Proton sequencer (Thermofisher, Waltham MA, USA). Comparison of the striatal transcriptome between *Thra*^*AMI/gn*^ mice and wild-type littermates was performed at PND7 and PND14 using the Ion AmpliSeq™ Transcriptome Mouse Gene Expression Kit (Thermofisher, Waltham MA, USA). All libraries were sequenced (> 10^7^ reads/library) on an Ion Proton sequencer (Thermofisher, Waltham MA, USA). For RNAseq, a count table was prepared using htseq-count (Galaxy Version 0.6.1galaxy3) (45). Differential gene expression analysis was performed with DEseq2 (Galaxy Version 2.1.8.3) (46) using the following thresholds: FDR < 0.05; p-adjusted value < 0.05; fold-change >2 or <0.5; expression > 10 reads per million. Hierarchical clustering was performed using the Cluster 3.0 R package (Euclidian distance, Ward’s algorithm). Chromatin immunoprecipitation for Chip-Seq analysis was performed as described (16). Libraries were prepared from the immunoprecipitated fraction and the input fraction as control, and sequenced on Ion Proton. 27.10^6^ reads were obtained for each library. Reads were mapped on the mouse genome (GRCm38/mm10 version) using Bowtie2 ((Galaxy Version 2.3.4.2). MACS2 (Galaxy Version 2.1.1.20160309.0) was used for peak calling. Peaks with a score inferior to 60 were filtered out, as well as peaks overlapping the “blacklist” of common artefacts (https://www.encodeproject.org/annotations/ENCSR636HFF/). De novo motif search was performed using MEME-ChIP (47).

## Acknowledgements

We thank Catherine Etter, Lucas Jacquin, Florent Delannoy and Victor Valcárcel, who contributed to histological studies, Tiphany Laurens, who contributed to behavioral analysis and Karine Gauthier, who performed *in vivo* experiments with propylthiouracil /TH treatments. We thank Nadine Aguilera, Marie Teixeira and the ANIRA-PBES facility for help in transgenesis and mouse breeding and Benjamin Gillet, Sandrine Hughes and the PSI platform of IGFL for deep DNA sequencing. This work was supported by a grant from the French Agence Nationale de la Recherche (Thyromut2 program; ANR-15-CE14-0011-01).

