## Supplementary material for "A pivotal genetic program controlled by thyroid hormone during the maturation of GABAergic neurons in mice": Suppl. info.

**This PDF file includes:**

Figs. S1 to S4
Tables S1 to S3
Captions for movies S1 to S2

**Other supplementary materials for this manuscript include the following:**

Movies S1 to S2
Dataset S1

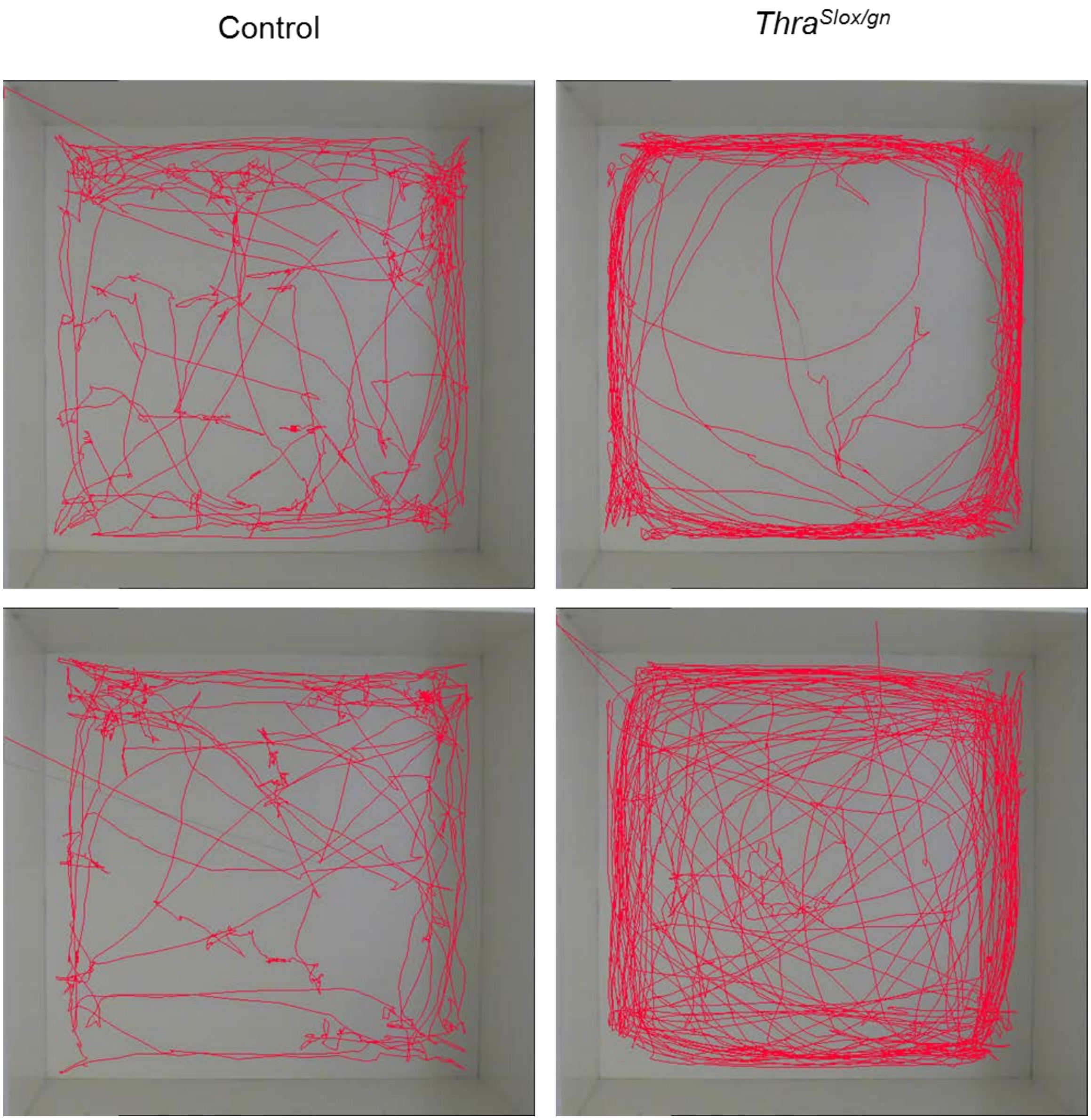


**Fig. S1** Representative tracks of adult control (left) and *Thra*$^{Slox/gn}$ (right) mice in a 5-min open-field test.

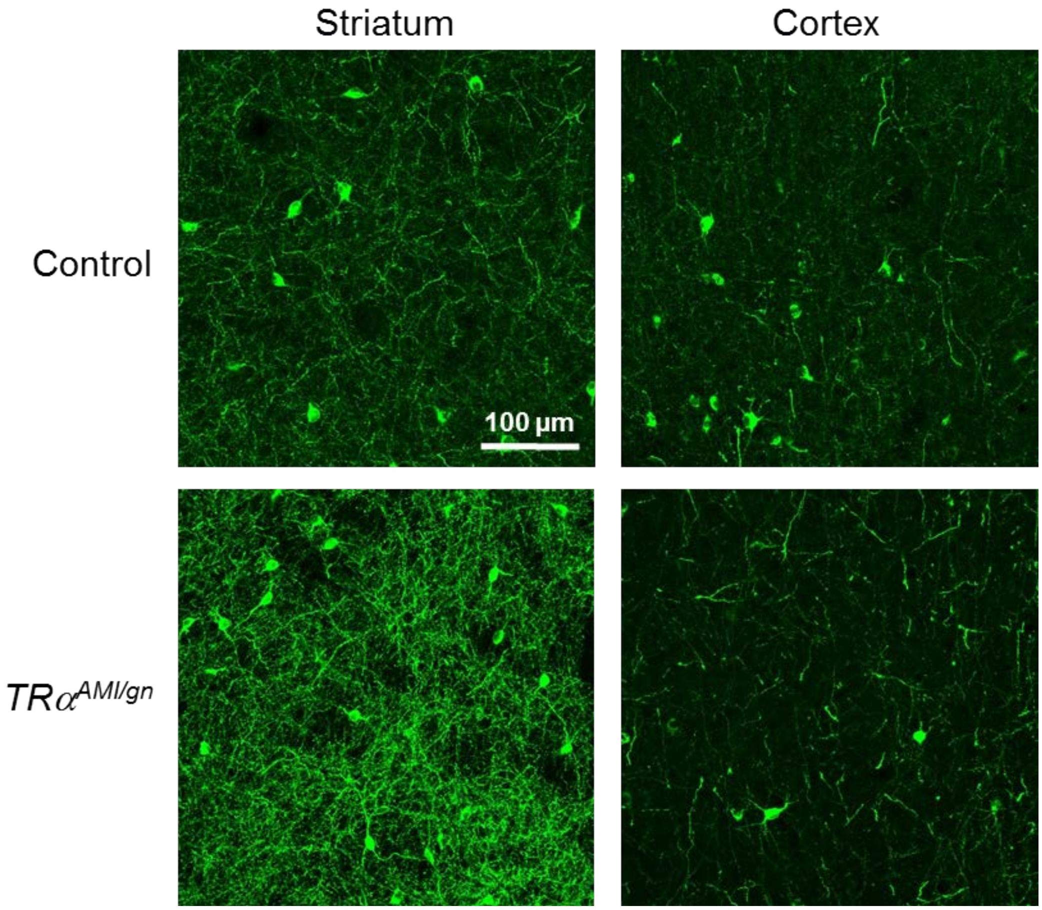


**Fig. S2.** Immunohistochemistry for neuropeptide Y in *Thra*$^{AMI/gn}$ and control mouse pups at PND14 in the striatum and cortex.

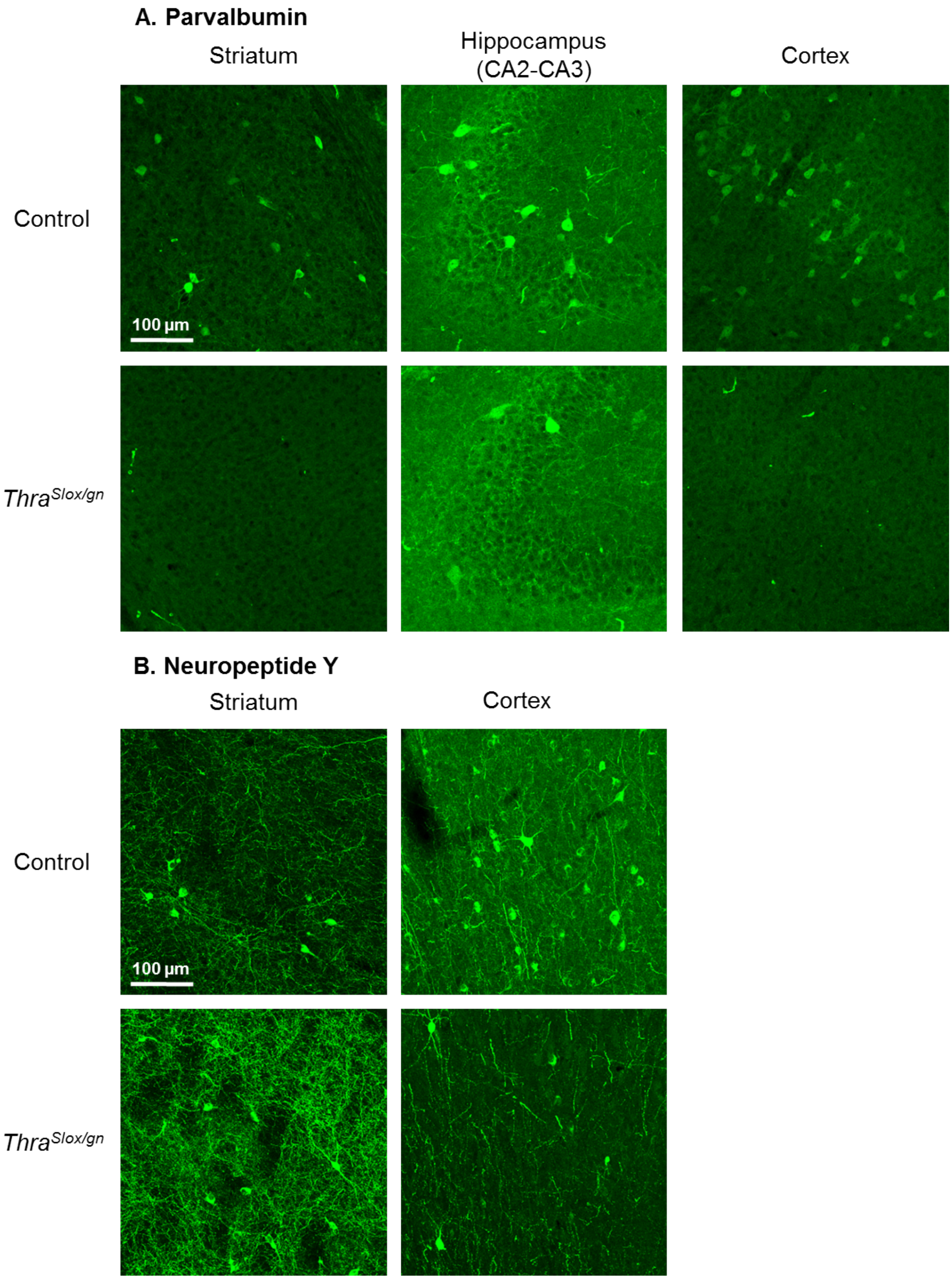


**Fig. S3.** Immunohistochemistry for parvalbumine (A) and neuropeptide Y(B) in *Thra*$^{Slox/gn}$ and control mouse pups at PND14 in selected brain regions.

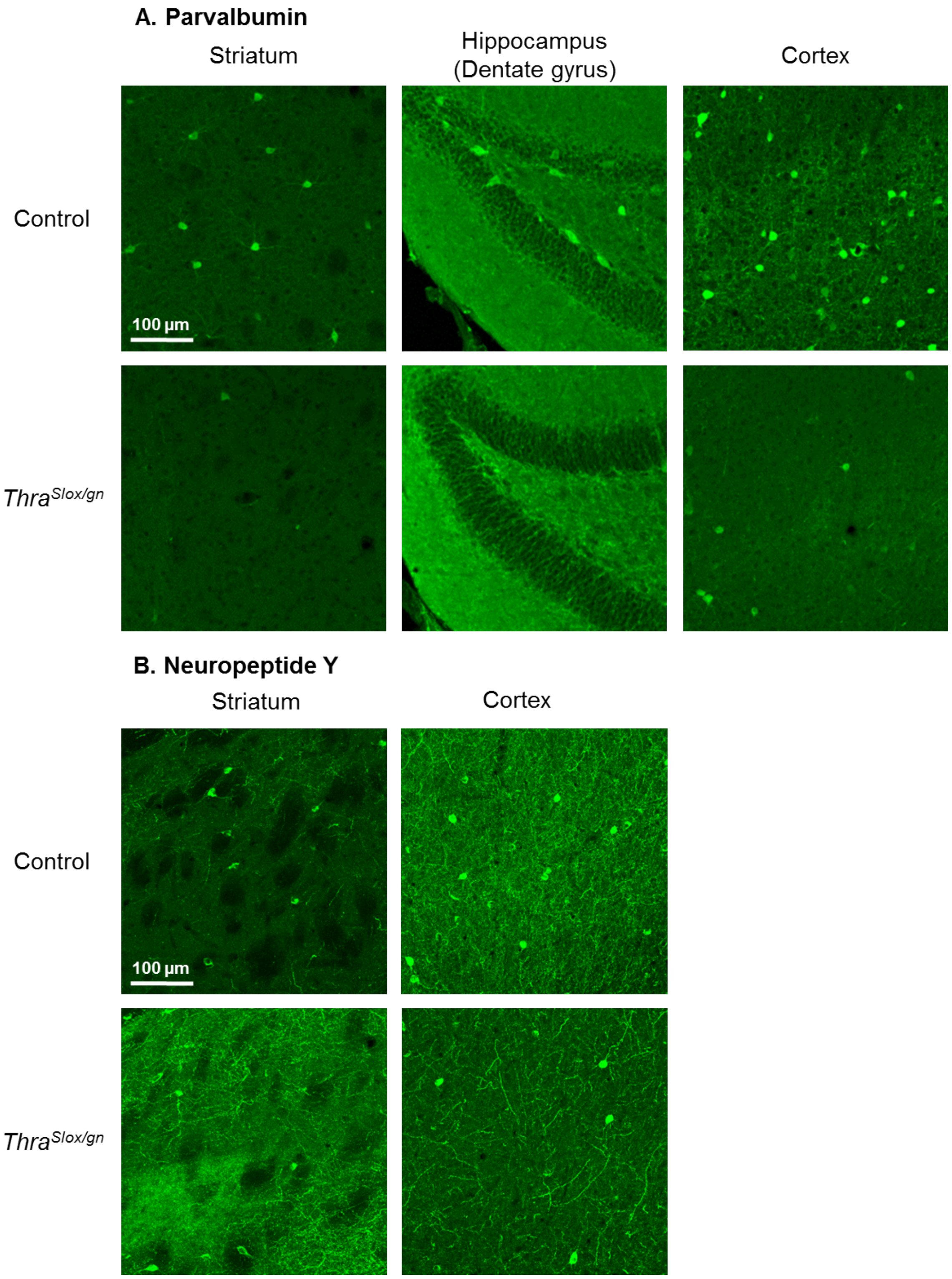


**Fig. S4.** Immunohistochemistry for parvalbumine (A) and neuropeptide Y(B) in adult *Thra*$^{Slox/gn}$ and control mice in selected brain regions.

**Table S1.** Relative abundance of several GABAergic neuron subtypes in *Thra*$^{AMI/gn}$, compared to control, mouse brains at PND14, as evidenced by immunohistochemistry.

| | | Cortex | | Hippocampus (DG) | | Hippocampus (CA) | | Striatum | |
|---|---|---|---|---|---|---|---|---|---|
| | | Control | *Thra* $^{AMI/gn}$ | Control | *Thra* $^{AMI/gn}$ | Control | *Thra* $^{AMI/gn}$ | Control | *Thra* $^{AMI/gn}$ |
| Parvalbumin neuronal density | Mean | 1.00 | 0.05[a] | 1.00 | 0.05[a] | 1.00 | 0.33[a] | 1.00 | 0.10[a] |
| | SD | 0.14 | 0.01 | 0.50 | 0.13 | 0.34 | 0.21 | 0.48 | 0.15 |
| | *n* | 11 | 10 | 11 | 12 | 11 | 10 | 10 | 8 |
| Neuropeptide Y neuronal density | Mean | 1.00 | 0.58[a] | nd | nd | nd | nd | 1.00 | 1.07 |
| | SD | 0.00 | 0.11 | nd | nd | nd | nd | 0.11 | 0.20 |
| | *n* | 5 | 6 | nd | nd | nd | nd | 5 | 6 |
| Neuropeptide Y fluorescence intensity | Mean | nd | nd | nd | nd | nd | nd | 1.00 | 1.31[a] |
| | SD | nd | nd | nd | nd | nd | nd | 0.06 | 0.25 |
| | *n* | nd | nd | nd | nd | nd | nd | 6 | 6 |
| Calretinin neuronal density | Mean | 1.00 | 0.75 | nd | nd | 1.00 | 4.90[a] | nd | nd |
| | SD | 0.12 | 0.54 | nd | nd | 0.41 | 4.10 | nd | nd |
| | *n* | 5 | 4 | nd | nd | 5 | 4 | nd | nd |
| Somatostatin neuronal density | Mean | 1.00 | 1.12 | 1.00 | 2.55[a] | 1.00 | 1.08 | 1.00 | 1.16 |
| | SD | 0.02 | 0.31 | 0.14 | 0.66 | 0.13 | 0.29 | 0.03 | 0.19 |
| | *n* | 6 | 5 | 6 | 5 | 6 | 5 | 6 | 5 |

[a] significantly different from control ($p$ < 0,05)

**Table S2.** Relative abundance of several GABAergic neuron subtypes in *Thra*$^{Slox/gn}$, compared to control, mouse brains at PND14 and in adults, as evidenced by immunohistochemistry.

| | | | Cortex | | Hippocampus (DG) | | Hippocampus (CA) | | Striatum | |
|---|---|---|---|---|---|---|---|---|---|---|
| | | | Control | *Thra* $^{Slox/gn}$ | Control | *Thra* $^{Slox/gn}$ | Control | *Thra* $^{Slox/gn}$ | Control | *Thra* $^{Slox/gn}$ |
| PND14 | Parvalbumin neuronal density | Mean | 1.00 | 0.01[a] | nd | nd | 1.00 | 0.56 | 1.00 | 0.10[a] |
| | | SD | 0.11 | 0.16 | nd | nd | 0.17 | 0.41 | 0.34 | 0.27 |
| | | *n* | 9 | 7 | nd | nd | 8 | 5 | 9 | 7 |
| | Neuropeptide Y neuronal density | Mean | 1.00 | 0.66[a] | nd | nd | nd | nd | 1.00 | 1.31 |
| | | SD | 0.25 | 0.26 | nd | nd | nd | nd | 0.17 | 0.25 |
| | | *n* | 6 | 6 | nd | nd | nd | nd | 6 | 6 |
| | Neuropeptide Y fluorescence intensity | Mean | nd | nd | nd | nd | nd | nd | 1.00 | 1.55[a] |
| | | SD | nd | nd | nd | nd | nd | nd | 0.14 | 0.15 |
| | | *n* | nd | nd | nd | nd | nd | nd | 6 | 6 |
| | Calretinin neuronal density | Mean | 1.00 | 1.23 | nd | nd | 1.00 | 2.30[a] | nd | nd |
| | | SD | 0.20 | 0.77 | nd | nd | 0.17 | 0.40 | nd | nd |
| | | *n* | 7 | 7 | nd | nd | 6 | 3 | nd | nd |
| Adult | Parvalbumin neuronal density | Mean | 1.00 | 0.31[a] | 1.00 | 0.13[a] | 1.00 | 0.50[a] | 1.00 | 0.39[a] |
| | | SD | 0.00 | 0.11 | 0.00 | 0.15 | 0.00 | 0.29 | 0.00 | 0.18 |
| | | *n* | 4 | 4 | 4 | 4 | 4 | 4 | 4 | 4 |
| | Neuropeptide Y neuronal density | Mean | 1.00 | 0.51[a] | nd | nd | nd | nd | 1.00 | 1.22 |
| | | SD | 0.00 | 0.12 | nd | nd | nd | nd | 0.00 | 0.31 |
| | | *n* | 4 | 4 | nd | nd | nd | nd | 4 | 4 |
| | Neuropeptide Y fluorescence intensity | Mean | nd | nd | nd | nd | nd | nd | 1.00 | 1.30[a] |
| | | SD | nd | nd | nd | nd | nd | nd | 0.00 | 0.10 |
| | | *n* | nd | nd | nd | nd | nd | nd | 5 | 5 |

[a] significantly different from control ($p$ < 0,05)

**Table S3.** List of 31 TRα1 direct target genes, as identified in the present study, using a combination of RNASeq and ChipSeq analyses.

| | FC *Thra* $^{AMI/gn}$ | FC hypoth. | FC TH response | KO mouse phenotype | Full name |
|---|---|---|---|---|---|
| Abcc12 | 0.44 | 0.49 | 2.08 | ND | ATP-binding cassette, sub-family C (CFTR/MRP), member 12 |
| Ace | 0.15 | 0.21 | 8.53 | Lethal | angiotensin I converting enzyme (peptidyl-dipeptidase A) 1 |
| Acvr1c | 0.19 | 0.44 | 2.46 | Normal | activin A receptor, type IC |
| Arg2 | 0.18 | 0.49 | 2.12 | Other | arginase type II |
| Ccm2 | 0.50 | 0.44 | 2.18 | Lethal | cerebral cavernous malformation 2 |
| Cd72 | 0.06 | 0.15 | 2.52 | Other | CD72 antigen |
| Cyp11a1 | 0.21 | 0.32 | 2.60 | Lethal | cytochrome P450, family 11, subfamily a, polypeptide 1 |
| Cyp2s1 | 0.06 | 0.11 | 16.51 | Normal | cytochrome P450, family 2, subfamily s, polypeptide 1 |
| Fblim1 | 0.27 | 0.50 | 2.44 | Other | filamin binding LIM protein 1 |
| Fgf16 | 0.18 | 0.34 | 3.69 | Lethal | fibroblast growth factor 16 |
| Fibcd1 | 0.44 | 0.28 | 2.00 | ND | fibrinogen C domain containing 1 |
| Gls2 | 0.28 | 0.44 | 3.09 | ND | glutaminase 2 (liver, mitochondrial) |
| Gpr139 | 0.27 | 0.12 | 3.80 | Other | G protein-coupled receptor 139 |
| Hr | 0.45 | 0.38 | 3.72 | Other | Hairless, lysine demethylase and nuclear receptor corepressor |
| Il17rc | 0.21 | 0.36 | 2.51 | Other | interleukin 17 receptor C |
| Kctd17 | 0.33 | 0.35 | 2.91 | Other | potassium channel tetramerisation domain containing 17 |
| Lpcat4 | 0.39 | 0.42 | 2.25 | Other | lysophosphatidylcholine acyltransferase 4 |
| Lynx1 | 0.46 | 0.30 | 2.00 | Neural Phenotype | Ly6/neurotoxin 1 |
| Me2 | 0.38 | 0.37 | 2.15 | Other | malic enzyme 2, NAD(+)-dependent, mitochondrial |
| Mme | 0.13 | 0.33 | 2.50 | Neural Phenotype | membrane metallo endopeptidase |
| Nrgn | 0.24 | 0.32 | 2.39 | Neural Phenotype | neurogranin |
| Nrtn | 0.29 | 0.44 | 2.89 | Other | neurturin |
| Pld5 | 0.38 | 0.33 | 2.35 | Normal | phospholipase D family, member 5 |
| Rasd2 | 0.40 | 0.39 | 2.32 | Neural Phenotype | RASD family, member 2 |
| Robo3 | 0.13 | 0.05 | 9.99 | Neural Phenotype | roundabout guidance receptor 3 |
| Sema7a | 0.19 | 0.28 | 2.30 | Neural Phenotype | sema domain, immunoglobulin domain (Ig), and GPI membrane anchor, |
| Sgpp2 | 0.16 | 0.15 | 3.18 | Other | sphingosine-1-phosphate phosphotase 2 |

**Table S3 (cont.).**

| | FC *Thra* $^{AMI/gn}$ | FC hypoth. | FC TH response | KO mouse phenotype | Full name |
|---|---|---|---|---|---|
| Shh | 0.29 | 0.39 | 2.52 | Neural Phenotype | sonic hedgehog |
| Slc22a3 | 0.35 | 0.34 | 2.13 | Neural Phenotype | solute carrier family 22 (organic cation transporter), member 3 |
| Slc26a10 | 0.22 | 0.31 | 2.40 | Neural Phenotype | solute carrier family 26, member 10 |
| Slc38a8 | 0.38 | 0.30 | 2.73 | ND | solute carrier family 38, member 8 |
| Slc41a1 | 0.45 | 0.39 | 2.12 | ND | solute carrier family 41, member 1 |
| Smpd3 | 0.44 | 0.48 | 2.02 | Other | sphingomyelin phosphodiesterase 3, |
| Tbc1d10c | 0.42 | 0.39 | 2.26 | Other | TBC1 domain family, member 10c |
| Ubash3b | 0.40 | 0.49 | 2.08 | Normal | ubiquitin associated and SH3 domain |

**Movie S1.** Representative video illustrating the occurrence of an epileptic seizure in a *Thra*$^{AMI/gn}$ mouse pup at postnatal day 17. The seizure occurred during the dark phase of the photoperiod and was video recorded using infrared lighting.

**Movie S2.** Representative video illustrating the occurrence of an epileptic seizure in a *Thra*$^{AMI/gn}$ mouse pup at postnatal day 13. Note the unsuccessful attempts of the mother to drag the pup back to the nest.
